# Human pancreatic β cell lncRNAs control cell-specific regulatory networks

**DOI:** 10.1101/096230

**Authors:** Ildem Akerman, Zhidong Tu, Anthony Beucher, Delphine M.Y. Rolando, Claire Sauty-Colace, Marion Benazra, Nikolina Nakic, Jialiang Yang, Huan Wang, Lorenzo Pasquali, Ignasi Moran, Javier Garcia-Hurtado, Natalia Castro, Roser Gonzalez-Franco, Andrew Stewart, Caroline Bonner, Lorenzo Piemonti, Thierry Berney, Leif Groop, Julie Kerr-Conte, Francois Pattou, Carmen Argmann, Eric Schadt, Philippe Ravassard, Jorge Ferrer

## Abstract

Recent studies have uncovered thousands of long non-coding RNAs (lncRNAs) in human pancreatic β cells. β cell lncRNAs are often cell type-specific, and exhibit dynamic regulation during differentiation or upon changing glucose concentrations. Although these features hint at a role of lncRNAs in β cell gene regulation and diabetes, the function of β cell lncRNAs remains largely unknown. In this study, we investigated the function of β cell-specific lncRNAs and transcription factors using transcript knockdowns and co-expression network analysis. This revealed lncRNAs that function in concert with transcription factors to regulate β cell-specific transcriptional networks. We further demonstrate that lncRNA *PLUTO* affects local three-dimensional chromatin structure and transcription of *PDX1,* encoding a key β cell transcription factor, and that both *PLUTO* and *PDX1* are downregulated in islets from donors with type 2 diabetes or impaired glucose tolerance. These results implicate lncRNAs in the regulation of β cell-specific transcription factor networks.

## Introduction

Transcriptome surveys have uncovered tens of thousands of mammalian transcripts longer than 200 nucleotides that have low protein-coding potential (Carninci et al., 2005; Derrien et al., 2012; Guttman et al., 2009). A small fraction of these long noncoding RNAs (lncRNAs) have been shown to control gene expression by modulating chromosomal structure, transcription, splicing, mRNA transport, stability or translation (Carrieri et al., 2012; Chen and Carmichael, 2009; Gong and Maquat, 2011; Lai et al., 2013; Luco and Misteli, 2011; Willingham et al., 2005; Yao et al., 2010). Specific lncRNAs have thus been implicated in various key processes, including random X chromosome inactivation, imprinting, the cell cycle, organogenesis, differentiation, pluripotency, and cancer progression (Guttman et al., 2011; Huarte et al., 2010; Hung et al., 2011; Klattenhoff et al., 2013; Kretz et al., 2013; Penny et al., 1996; Schmitt and Chang, 2013; Sleutels et al., 2002; Ulitsky et al., 2011). Despite these wide ranging biological roles, the fraction of lncRNAs that is genuinely functional, and the true impact of lncRNAs in human biology and disease remains poorly understood.

Pancreatic β cells regulate glucose homeostasis by secreting the insulin, and play a central role in the pathogenesis of major forms of diabetes mellitus. Recently, more than 1100 lncRNAs were identified in human pancreatic islets and purified β cells (Moran et al., 2012), as well as in mouse pancreatic islet cells (Benner et al., 2014; Ku et al., 2012; Moran et al., 2012). A large fraction of human β cell lncRNAs are cell-specific, and several are known to be activated during β cell differentiation (Moran et al., 2012). This cellular specificity has also been noted for lncRNAs in other cell types (Cabili et al., 2011; Derrien et al., 2012), and points to the possibility that lncRNAs may regulate genetic programs important for lineage-specific differentiation or specialized cellular functions. Further, several β cell lncRNAs were shown to be regulated by extracellular glucose concentrations, suggesting a potential role of lncRNAs in the functional adaptation of β cells to increased insulin secretory demands (Moran et al., 2012). Some islet lncRNAs map to loci that contain polygenic or Mendelian defects associated with human diabetes, while selected lncRNAs show deregulation in islets from organ donors with human type 2 diabetes (T2D) (Fadista et al., 2014; Moran et al., 2012). Collectively, these properties define a newly identified class of candidate regulators of β cell differentiation and function, with potential implications for human diabetes mellitus. However, the true relevance of β cell lncRNAs depends on whether they elicit a physiological function in human β cells, which remains to be addressed systematically.

In the current study, we have focused on a set of lncRNAs that show restricted expression in human pancreatic β cells, and have tested the hypothesis that they regulate β cell gene expression. Our studies have uncovered a regulatory network in which lineage-specific lncRNAs and transcription factors (TFs) control common genes. Furthermore, we show that lncRNAs frequently regulate genes associated with clusters of islet enhancers, which have previously been shown to be the primary functional targets of islet-specific TFs. Detailed analysis of a specific lncRNA named *PLUTO* controls *PDX1*, a master regulator of pancreas development and β cell differentiation, and thereby modulates the *PDX1*-dependent transcriptional program. Finally, we show that *PLUTO* and *PDX1* are downregulated in islets from organ donors with type 2 diabetes or impaired glucose tolerance, suggesting a potential role in human diabetes.

## Results

### Human β cell IncRNA knockdowns cause profound transcriptional phenotypes

To directly test the regulatory function of pancreatic β cell lncRNAs, we carried out loss of function experiments in a glucose-responsive human islet β cell line, EndoC-ßH1 (Ravassard et al., 2011). We chose a human model because only some human lncRNAs are evolutionary conserved (Derrien et al., 2012; Moran et al., 2012; Okazaki et al., 2002; Pang et al., 2006), and we perturbed the function of lncRNAs through RNAi-based transcript knockdowns rather than genomic deletions because deletions could potentially disrupt cis-regulatory elements. We thus designed lentiviral vectors that contain RNA Polymerase II-transcribed artificial miRNAs (hereafter referred to as amiRNA) with perfect homology to the target sequence so as to elicit target cleavage. The amiRNAs contain an artificial stem sequence targeting our lncRNA of choice as well as flanking and loop sequences from an endogenous miRNA to allow their processing as pre-miRNA by the RNAi pathway (**Figure S1A**). As a reference, we used the same strategy to knockdown TFs that are well known to regulate gene expression in pancreatic islets, as well as five different non-targeting amiRNA sequences as controls.

The lncRNAs selected for knockdown were derived from a shortlist of 25 lncRNAs that showed (i) a markedly enriched expression in human islets and FACS-purified β cells relative to exocrine pancreas and a panel of non-pancreatic tissues, (ii) expression in the EndoC-βH1 β cell line, and (iii) a chromatin profile in human islets that was consistent with an active promoter (**Figure S1C-D**). Of these 25 lncRNAs, 12 were shortlisted because they were near a protein-coding gene that has an important function in β cells. The lncRNAs had variable subcellular enrichment patterns (**Figure S1B**) and eight of the 12 lncRNAs had detectable transcripts in orthologous or syntenic mouse regions (Table S1)(Moran et al., 2012). We then screened four amiRNA sequences for each of the 12 lncRNAs and identified two efficient (>50% knockdown) amiRNAs for 7 lncRNAs, and one efficient amiRNA sequence for the other five lncRNAs (**Figure S1E**). Two efficient amiRNAs were also obtained for five essential islet TFs (*HNF1A, GLIS3, MAFB, NKX2.2, PDX1*). We thus transduced EndoC-βH1 cells with lentiviruses expressing each amiRNA. This was done in duplicate, or in triplicate for lncRNAs that only had one efficient amiRNA. 80 hrs post-transduction, RNA was harvested and hybridized to oligonucleotide microarrays (**Figure 1A**). For each target gene, we combined expression data from all knockdowns and compared them to the control transductions with five different control amiRNAs to identify genes that were differentially expressed at a significance level of p<10^−3^ (ANOVA) (**Figure 1B**).

**Figure 1.**
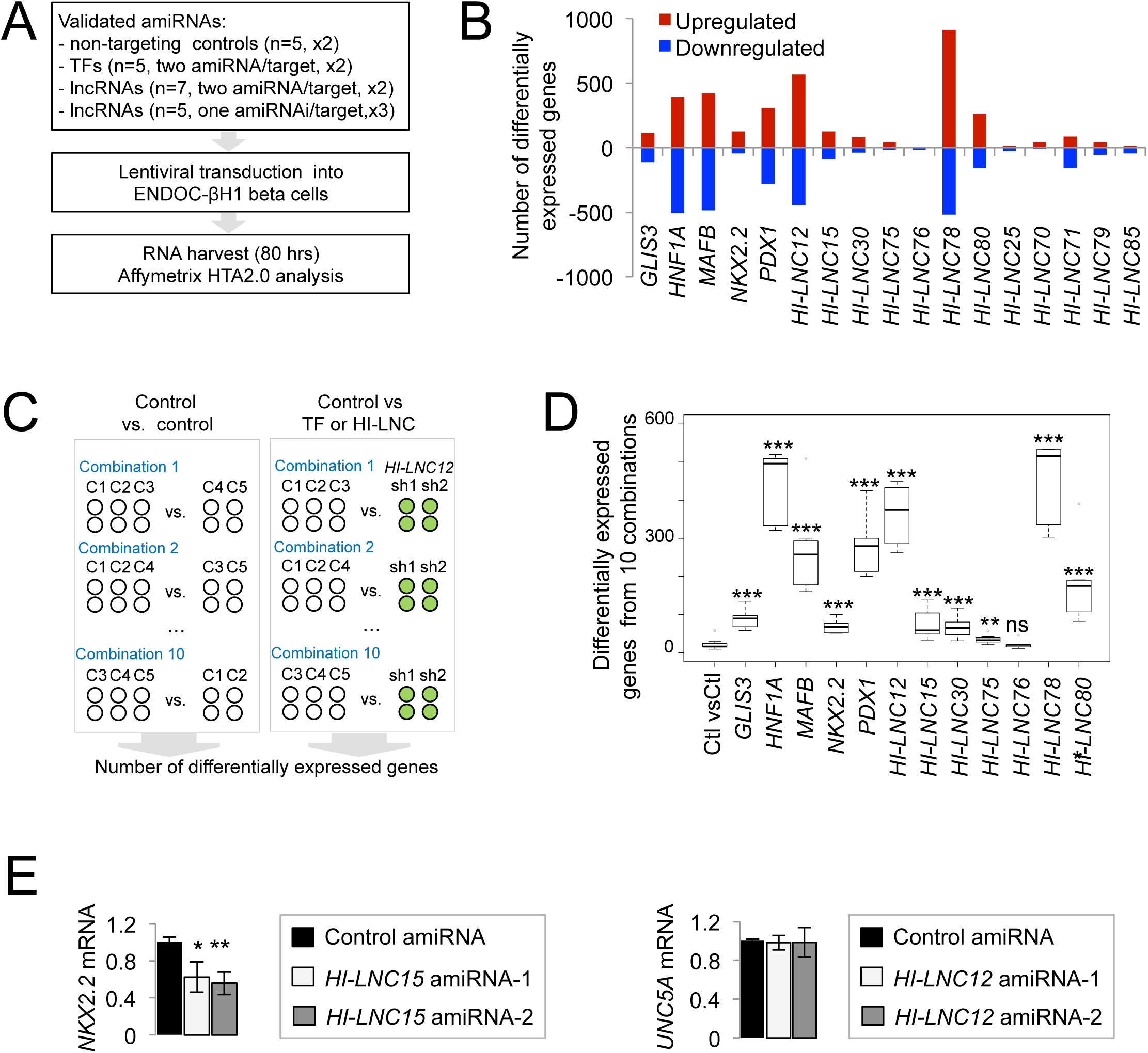
Knockdown of selected β cell lncRNAs leads to transcriptional phenotypes. (A) Schematic representation of the experimental plan. Lentiviral encoded amiRNAs were validated and transduced in duplicate (x 2) or triplicate (x 3) into ENDOC-βH1 cells as indicated, and then analyzed with oligonucleotide expression arrays. (B) Differential gene expression analysis revealed genes that show significant up or downregulation after knockdown of TFs or lncRNAs. For each TF or lncRNA, we combined all replicates transduced with the different target-specific amiRNAs, and compared these to all replicates from 5 non-targeting controls. Differential expression was determined at p<10^−3^ (ANOVA). (C) We compared gene expression data from all 10 possible combinations of 3 vs. 2 control non-targeting amiRNAs. Similarly, the two independent amiRNAs that target each TF or lncRNA were compared to all 10 possible combinations of 3 control amiRNAs. For this analysis we only considered the 7 lncRNAs that were targeted by two independent amiRNAs. (D) Control comparisons result in a low number of differentially regulated genes (average 15 genes), while most TF and lncRNA comparisons yield higher numbers of differentially regulated genes. ***p<10^−4^, **p<0.01, ns: not significant, as compared to control comparisons, Mann-Whitney test. (E) *HI-LNC15* regulates its neighboring gene *NKX2.2,* while *HI-LNC12* knockdown (KD) does not affect its adjacent active gene, *UNC5A* (left panel). Further examples are shown in Figure S1G. RNAs were normalized to *TBP* mRNA and expressed relative to control amiRNAs; n=3, error bars represent SEM, **p<0.01, *p<0.05 (Students t-test).

As expected, the knockdown of islet TFs consistently produced transcriptional phenotypes (**Figure 1B**). Remarkably, the knockdown of 9 of the 12 islet lncRNAs also caused transcriptional changes (**Figure 1B**, **S1F**). A more detailed analysis showed that some of the lncRNAs that presented knockdown phenotypes had visible effects on a neighboring gene, suggesting a possible cis-regulatory mechanism, although other such lncRNAs did not appear to affect neighboring genes, and may thus function through trans-regulatory mechanisms (**Figure 1E** and **S1G**). These loss of function experiments with selected lncRNAs therefore suggested that lncRNAs can regulate the expression of pancreatic β cell genes.

Gene silencing using the RNAi pathway can theoretically lead to nonspecific gene deregulation. In our experimental model, a significant nonspecific result would occur if two unrelated amiRNAs elicited changes in a common set of genes that were not observed in the panel of control non-targeting amiRNAs. To assess the likelihood that two unrelated amiRNA sequences elicit such an effect, we studied the 5 sets of control (non-targeting) amiRNAs, compared all 10 possible combinations of 2 vs. 3 control amiRNAs, and determined the number of differentially expressed genes (**Figure 1C**). Likewise, for each TF or lncRNA which had two valid amiRNAs, we compared the two target-specific amiRNAs against all possible combinations of three control amiRNAs (**Figure 1C**). As seen in **Figure 1D**, control vs. control comparisons generated a median of 16 (IQR=15-22) differentially expressed genes, whereas all five TFs and six of the seven lncRNA knockdowns led to a significantly higher number of differentially expressed genes (Mann-Whitney test p<10^−4^ for all lncRNA/TF vs. control comparisons except *HI-LNC75,* p=0.004, and *HI-LNC76,* p>0.5). These results show that the observed phenotypes are unlikely to be caused by unspecific effects of amiRNAs, and indicate that the sequence-specific inhibition of selected islet lncRNAs can result in transcriptional changes comparable in magnitude to the inhibition of well established islet transcriptional regulators.

The primary function of β cells is to synthesize and secrete insulin in response to changes in glucose concentrations. Amongst the genes that showed functional dependence on lncRNAs we identified numerous genes that are known to regulate transcription or secretion in β cells, including *RFX6, PDX1, CACNA1D, ATP2A3, ROBO1* and 2, *PDE8A, ATP6AP1, KCNJ15, TRPM3, ERO1LB* and *HADH* (**Figure 2A**) (Anderson et al., 2011; Li et al., 2010; Louagie et al., 2008; Okamoto et al.,2012; Smith et al., 2010; Tian et al., 2012; Varadi and Rutter, 2002; Wagner et al., 2008; Yang et al., 2013; Zito et al., 2010). We therefore measured insulin content and glucose-stimulated insulin secretion (GSIS) in T-antigen excised EndoC-βH3 cells after knocking down four lncRNAs that showed the strongest transcriptional phenotypes *(HI-LNC12, HI-LNC78, HI-LNC80* and *HI-LNC71)*. Congruent with the broad transcriptional phenotype, we observed reduced insulin content and consequently impaired glucose-stimulated insulin secretion for *HI-LNC12, HI-LNC78*, and *HI-LNC71* knockdowns (**Figure 2B**). For *HI-LNC78,* a glucose-regulated islet transcript (Moran et al, 2012) that is orthologous to mouse *Tunar* and zebrafish *megamind^(linc-birc6)^* lncRNAs (Ulitsky et al., 2011), there was a reduction in GSIS after correcting for the reduction in insulin content (p=0.002) (**Figure S2A**). To further validate these effects, the same lncRNAs were downregulated using antisense locked nucleic acids (LNA^TM^ GapmeRs, Exiqon), which also led to impaired insulin secretion after knockdown of *HI-LNC12* and *HI-LNC78* (**Figure S2B**). Taken together, lncRNA knockdown studies identified lncRNAs that modulate gene expression and consequently insulin secretion in a human β cell line.

**Figure 2.**
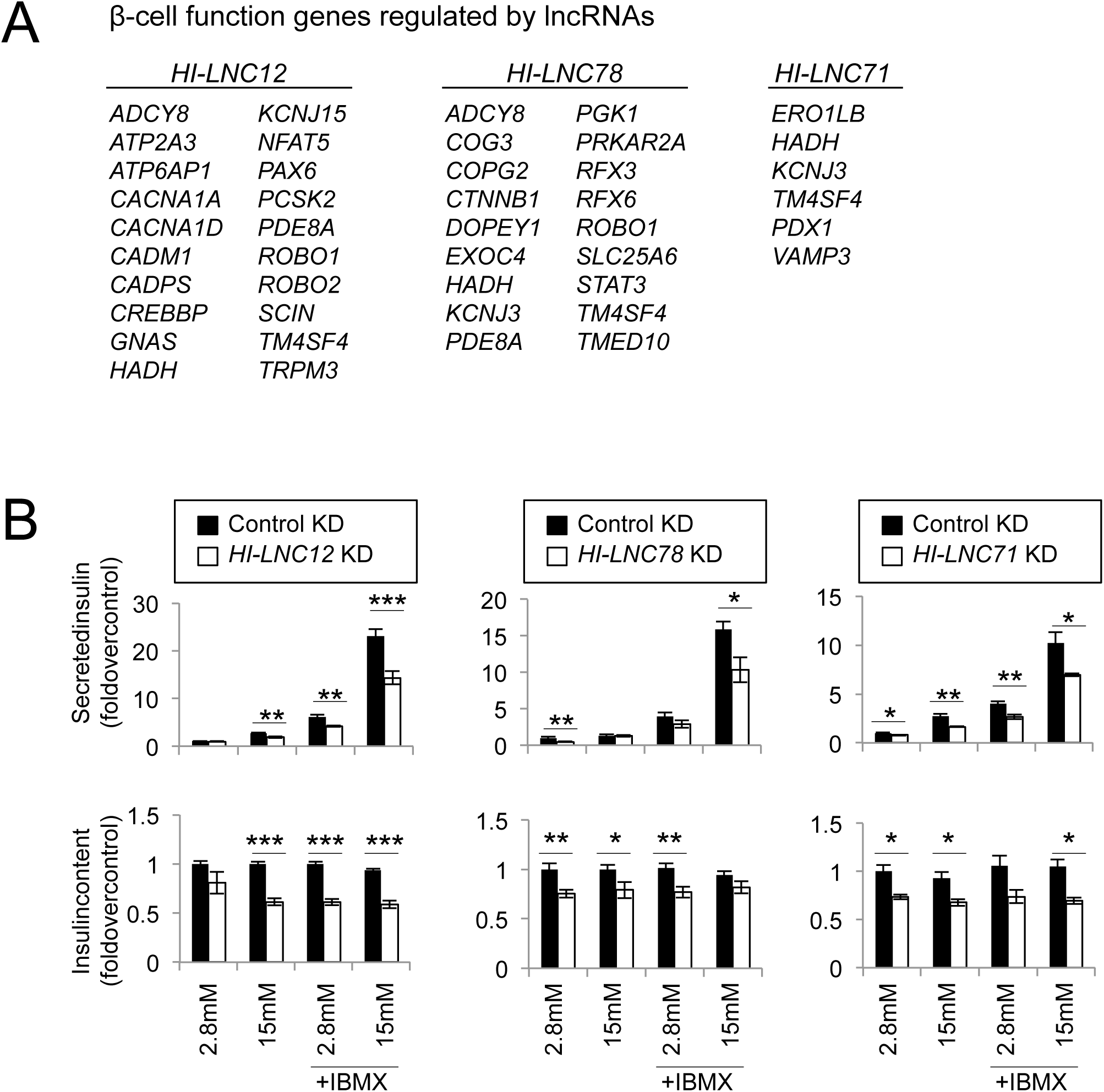
Knockdown of lncRNAs impairs insulin secretion. (A) Examples of genes known to play a role in β cell function regulated by islet lncRNAs. (B) Glucose-stimulated insulin secretion was tested on T-antigen excised EndoC-βH3 cells after transduction with amiRNAs targeting indicated lncRNAs or controls. Secreted or total insulin content was normalized to the number of cells per well and expressed as fold change over control amiRNA treatment at 2.8 mM glucose. Each bar represents an average from two independent amiRNA vectors and 12 separate wells, from two independent experiments. Error bars represent SEM, *** p<10^−3^, ** p<0.01, *p*0.05 (Student’s t-test).

### Human islet lncRNAs and TFs regulate common gene expression programs

To gain insight into the expression programs that are regulated by islet-specific lncRNAs and TFs, we compared their knockdown gene expression phenotypes. We first assessed changes in gene expression occurring after knockdown of the different islet TFs, and found high Pearson correlation values for all pairwise comparisons (r = 0.4-0.8, p<10^−27^)(**Figure 3A**, **S3**). This finding is consistent with the notion that islet-specific TFs often bind to common genomic targets and function in a combinatorial manner (Pasquali et al., 2014; Qiu et al., 2002; Wilson et al., 2003). Interestingly, the transcriptional changes that occurred after the inhibition of several lncRNAs significantly correlated with those observed following inhibition of TFs (**Figure 3A** and **S3**, see also a cluster analysis of TF and lncRNA-dependent changes in **Figure 3B**). Some pairwise comparisons that illustrate this finding include *HI-LNC78-* dependent gene expression changes, which correlated highly with *HNF1A* and *MAFB* dependent changes (Pearson’s r=0.87 and 0.89, respectively, p<10), and *HI-LNC15-dependent* changes, which correlated with those occurring after knockdown of *NKX2-2* (r=0.67, p=10^−32^) (**Figure 3C**). The results from these gene knockdown experiments therefore indicate that selected islet-specific lncRNAs and TFs can regulate common gene expression programs.

**Figure 3.**
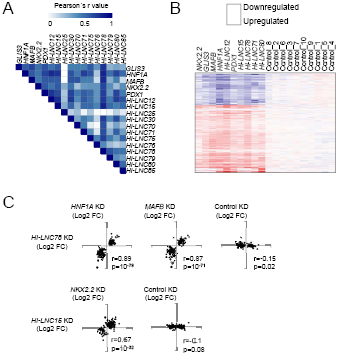
Human islet TFs and lncRNAs regulate common genes. (A) Heatmap displaying Pearson r values for all pairwise comparisons of fold-changes in gene expression after knockdown of TFs and lncRNAs. Only genes significantly dysregulated at p<10^−3^ in at least one condition were included in the analysis. (B) Unsupervised clustering analysis of fold-change values after knockdown of 5 TFs and the 5 lncRNAs that displayed the strongest transcriptional changes. Only genes that were dysregulated at p<10^−3^ in at least one knockdown were selected. Blue represents downregulated and red represents upregulated genes. Controls represent control comparisons as described for **Figure 1**. (C)Examples of highly correlated transcriptional phenotypes. The plots show fold-change values (Log2) after knockdown of indicated pairs of genes. Only the top 100 most regulated genes for any of the two knockdowns were plotted. Pearson’s correlation (r) and p-values are displayed.

### Islet TFs and lncRNAs co-regulate genes associated with enhancer clusters

Recent studies have revealed that islet TFs regulate cell-specific transcription by targeting clusters of enhancers, and in particular clusters with enhancers that are bound by multiple islet TFs (Pasquali et al., 2014). Enhancer clusters share many features with regulatory domains that have otherwise been defined as “stretch enhancers” or “super-enhancers” (Pasquali et al., 2014; Pott and Lieb, 2015). Given that knock-down of islet lncRNAs and TFs suggested that they regulate similar genes, we asked if islet lncRNAs also regulate enhancer cluster-associated genes. As expected, Gene Set Enrichment Analysis (GSEA) showed that genes with islet-enriched expression, genes associated with enhancer clusters, or genes associated with enhancers that are bound by multiple TFs were downregulated after knockdown of all five TFs, whereas this was not observed for ten control sets of genes expressed at similar levels (**Figure 4**, **Figure S4A,B**). Likewise, genes associated with enhancer clusters and those showing islet-specific expression were also enriched among genes that were downregulated after knockdown of *HI-LNC12, 15, 30, 78, 80, 85* and *71* (**Figure 4**, **Figure S4A,B**). These results therefore indicate that islet-specific TFs and lncRNAs often co-regulate genes that are associated with enhancer clusters.

**Figure 4.**
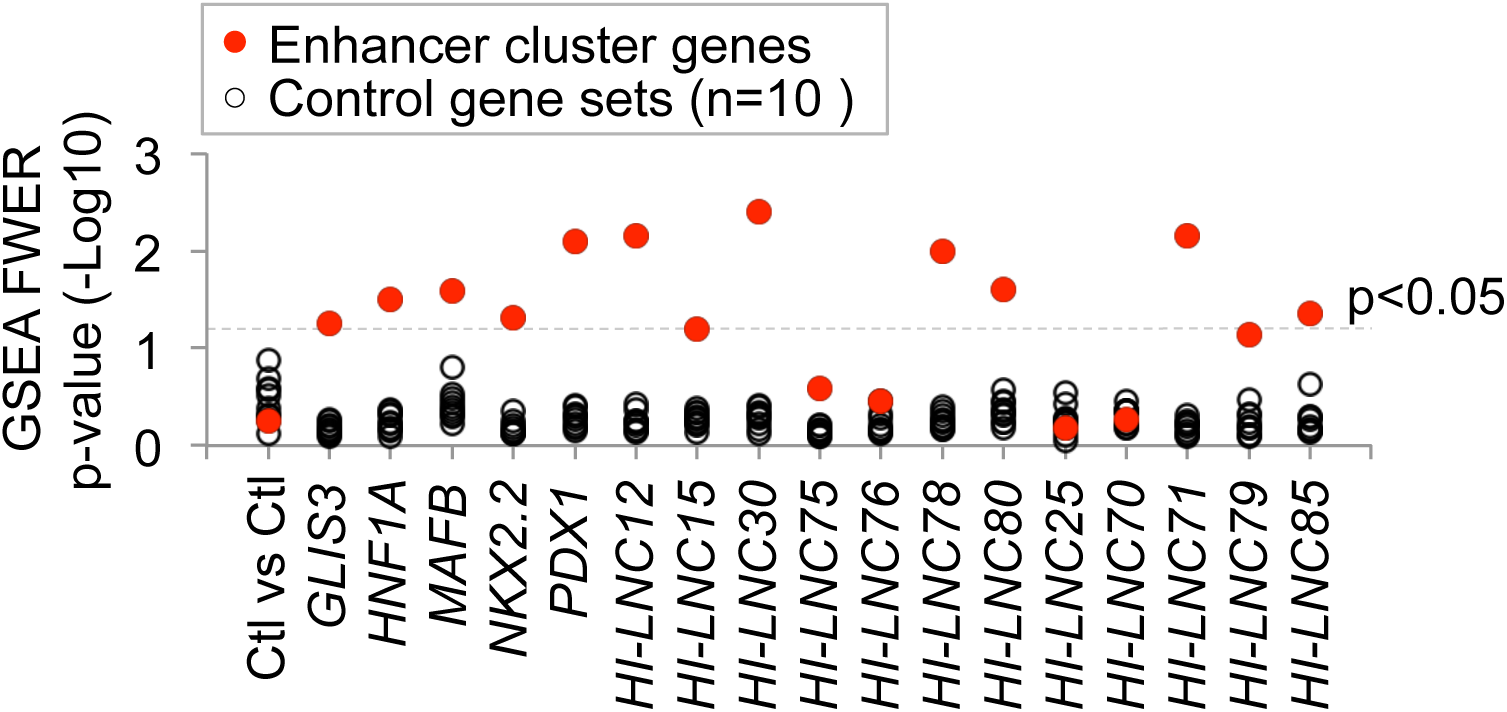
LncRNAs regulate enhancer cluster genes. Gene Set Enrichment analysis (GSEA) showed that genes that were downregulated upon knockdown of either islet TFs or lncRNAs were enriched in a set of 694 genes that is associated with human islet enhancer clusters (red dots), but not in 10 control gene sets (black dots) that were expressed at similar levels as enhancer cluster genes.

### β cell lncRNAs and TFs form part of islet-specific co-expression networks

We next used an independent experimental approach to validate the observations that human β cell lncRNAs and TFs regulate common gene expression programs. This involved the analysis of gene modules that show co-expression across a panel of human islet RNA samples. Analogous approaches have been employed to reveal sets of genes that share functional relationships (Derry et al., 2010; Kim et al., 2001; Pandey et al., 2010; Segal et al., 2003; Stuart et al., 2003; Su et al., 2011). We implemented this analysis using weighted gene co-expression analysis (WGCNA) of RNA-seq profiles from 64 human pancreatic islet samples. This identified 25 major gene modules containing >100 genes, named M1-M25, which showed highly significant co-expression across human islet samples (**Figure 5A**, **Table S2**). We next determined which co-expression modules contained islet lncRNAs. Rather than using our previously defined set of lncRNAs, this analysis was performed with a set of 2373 β cell lncRNAs that was newly annotated using ∼5 billion stranded RNA-seq reads pooled from 41 islet samples (**Table S3, Figure S5A**). β cell lncRNAs were found to be enriched in seven pancreatic islet co-expression modules (M3, M7, M12, M13, M18, M20, M21) (**Figure 5B**).

**Figure 5.**
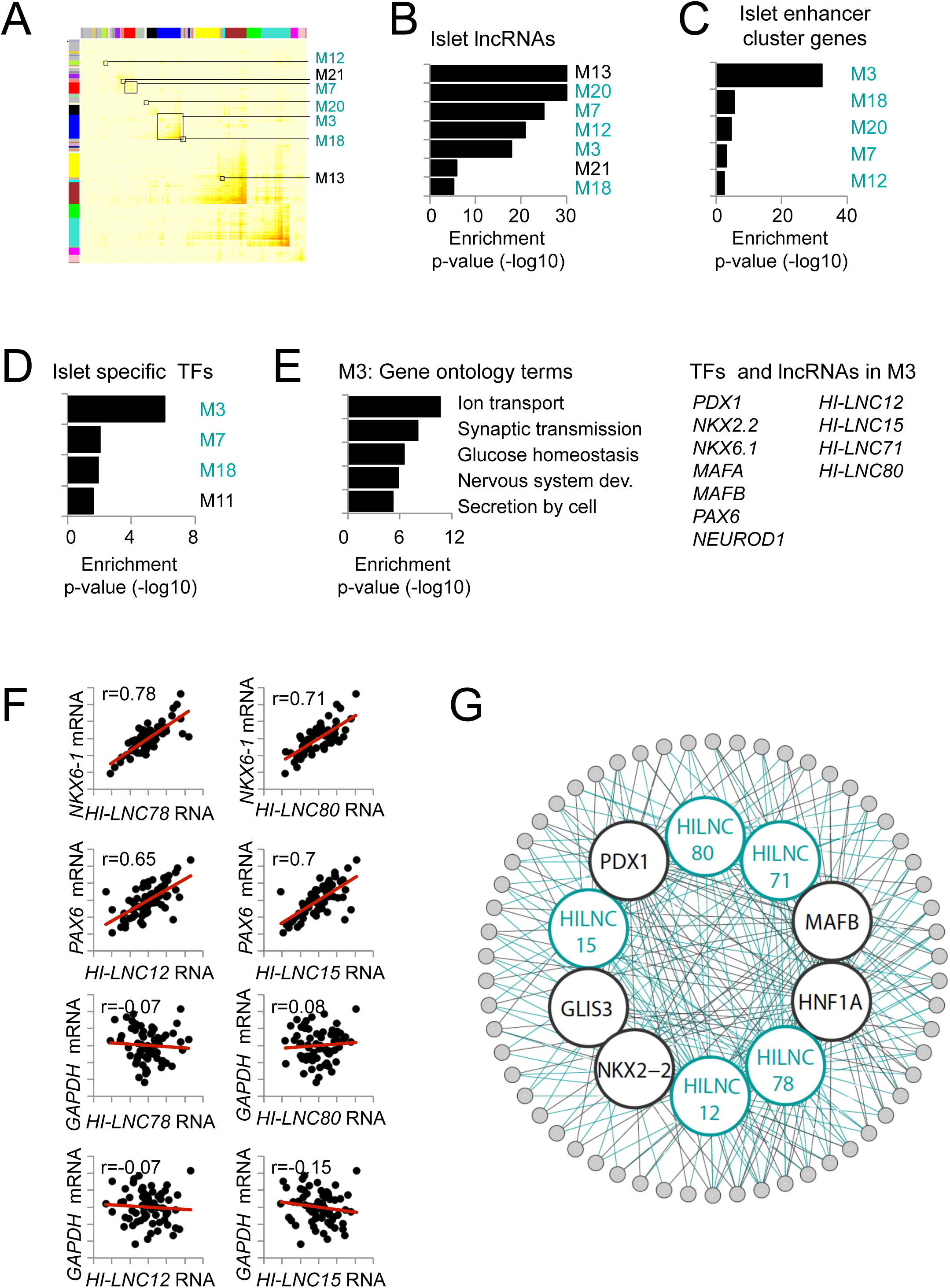
Islet-specific coding and noncoding RNAs form shared co-expression modules. (A) Topological overlap matrix representing co-expression modules that were co-regulated across 64 human islet samples. Modules that were enriched in lncRNAs are marked with squares (hypergeometric test, p<10^−2^). (B-D) Co-expression modules that showed enrichment in (B) islet lncRNAs, (C) islet enhancer cluster (EC)-associated genes, or (D) a set of 94 islet-enriched TF genes. Five modules (M3, M7, M12, M18 and M20, marked in blue) out of seven modules that were enriched in lncRNAs were also enriched in ECs and TFs. (E) Module M3 was enriched in typical islet-specific biological process annotations. The right panel shows examples of islet TFs and lncRNAs in module M3. (F) Correlation of indicated lncRNAs and β cell-specific TF mRNAs across 64 islet samples. *GAPDH* is shown as a non β cell reference. Pearson’s correlation values are displayed in the top left corner. The axes show expression values normalized across 64 islet samples. (G) Network diagram illustrating that TFs and lncRNAs often co-regulate the same genes, many of which were associated with enhancer clusters.

We next characterized the nature of these seven lncRNA-enriched co-expression modules. Five of these (M3, M7, M12, M18, M20) were enriched in genes associated with pancreatic islet enhancer clusters (**Figure 5A-C**, marked in blue). Two other modules (M13, M21) were enriched for ubiquitously expressed genes involved in mRNA translation and metabolic pathways (**Figure S5B**). Amongst the modules enriched in lncRNAs and enhancer clusters, three (M3, M7, M18) were also enriched in islet-specific TF genes (**Figure 5D**), and two of these modules (M3, M7) contained nine of the 12 lncRNAs that had been knocked down in EndoC-βH1 cells. Module M3, the largest of the seven lncRNA-enriched modules, featured gene ontology (GO) terms associated with prototypical islet cell functions and contained several islet TFs and lncRNAs (**Figure 5E**). In keeping with these findings, we found numerous instances of islet lncRNAs and known cell-specific TFs that showed a tight correlation of gene expression levels across human islet samples (**Figure 5F**, **S5C**). These findings thus indicated that β cell-specific lncRNAs, TFs, and genes associated with islet enhancer clusters form part of common expression programs.

Further analysis is consistent with the notion that lncRNAs play a functional role in driving gene expression variation in the lncRNA-enriched co-expression modules. First, the subset of lncRNAs that were shown to regulate an adjacent gene in knockdown studies also exhibited a particular high co-regulation with the adjacent gene across islet samples (**Figure S1G**). This observation was extended to define 292 lncRNAs that displayed a highly significant (p<10^−7^) correlation of expression with an adjacent protein-coding gene in the panel of human islet samples, and are thus candidate cis-regulatory lncRNAs (**Table S6**). Second, we analyzed all genes that were significantly downregulated in EndoC-βH1 cells after knocking down *HI-LNC12, 71, 78* and *80,* and found that they were also enriched amongst genes in human islet modules M3, M7 and M18, but not in size-controlled modules (**Figure S5D**). In summary, co-expression analysis of native human islets corroborated the findings observed with amiRNA-based perturbations in EndoC-βH1 cells, and indicated that a group of islet lncRNAs and TFs form part of common transcriptional networks that target clusters of pancreatic islet enhancers (**Figure 5G**).

### Deregulation of β cell lncRNAs in human T2D

The identification of functional lncRNAs led us to explore whether some lncRNAs are abnormally expressed in human T2D, and might thus be relevant to the pathogenesis of this disease. We therefore analyzed our new set of 2373 lncRNAs in a recently reported gene expression dataset that includes human islet samples from donors diagnosed with T2D or impaired glucose tolerance (IGT) (Fadista et al., 2014). Our results showed that, despite the fact that gene expression across human islet donors is highly variable, the expression of 15 and 100 lncRNAs was significantly altered in islets from T2D and IGT vs. non-diabetic donors, respectively (adjusted p<0.05) (**Figure S6A**, see **Table S7** for a complete list). This finding suggests a potential role of functional β cell lncRNAs in driving some of the β cell gene expression changes that are associated with T2D.

### *PLUTO* regulates *PDX1*, an essential transcriptional regulator

To explore how β cell lncRNAs can regulate cell-specific transcriptional networks, we focused on *HI-LNC71,* a nuclear-enriched transcript (**Figure S1B**) that is transcribed from a promoter that is located ∼3 kb upstream of *PDX1*, in an antisense orientation (**Figure S6B**). PDX1 is an essential transcriptional regulator of pancreas development and β cell function that has been implicated in genetic mechanisms underlying Mendelian and type 2 diabetes (Ahlgren et al., 1998; Jonsson et al., 1994; Offield et al., 1996; Stoffers et al., 1997). Based on this genomic location, we renamed *HI-LNC71* as *PLUTO*, for *<u>P</u>DX1* <u>L</u>ocus <u>U</u>pstream <u>T</u>ranscript.

The potential importance of *PLUTO* was strengthened by the observation that *PLUTO* was among the most markedly downregulated lncRNAs in islets from T2D or IGT donors (adjusted p-value = 0.07 and 0.005, respectively, **Figure 6A**, **S6B**). Interestingly, PDX1 was also downregulated in islets from donors with T2D and IGT (**Figure 6A**).

**Figure 6.**
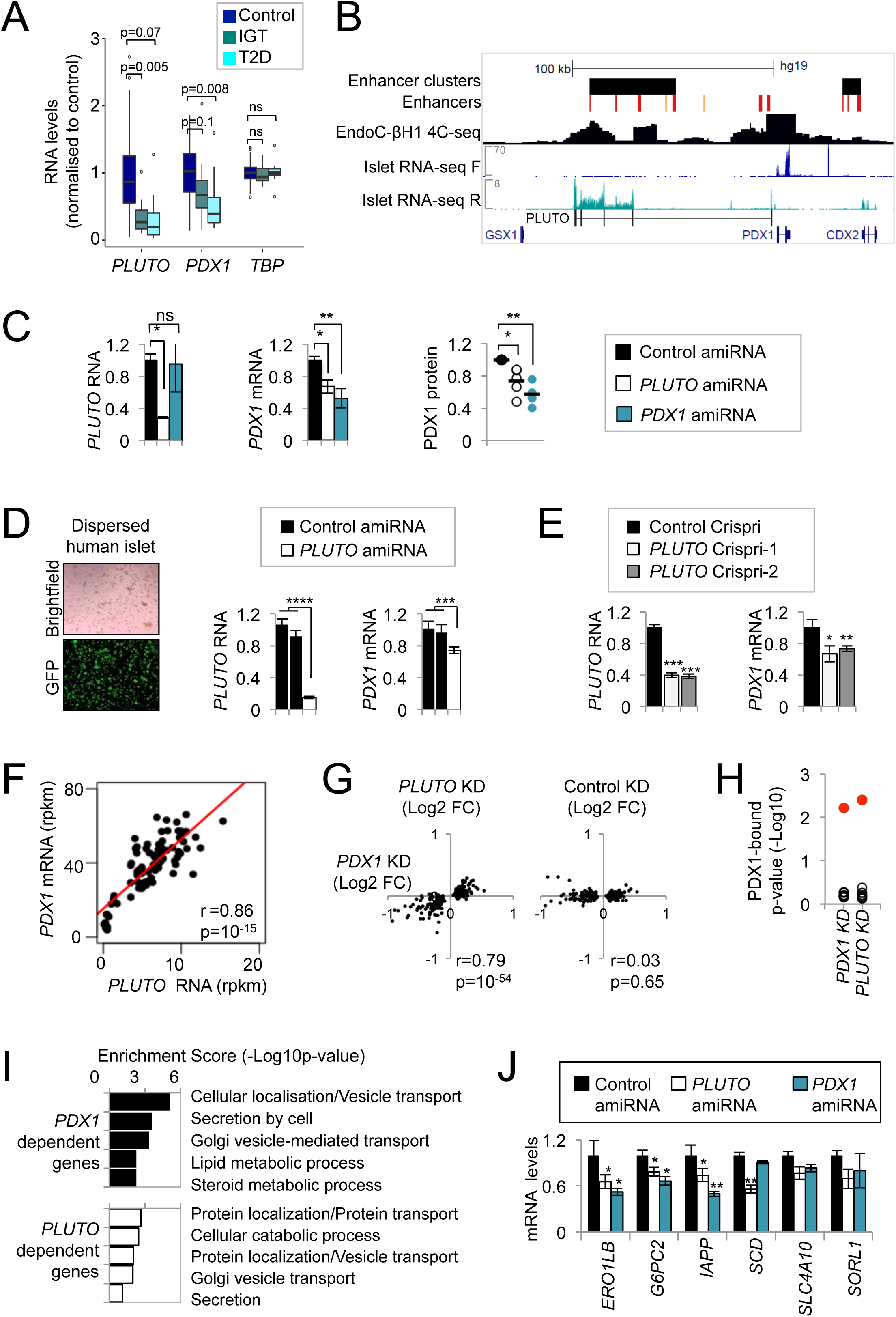
*PLUTO* knockdown decreases *PDX1* mRNA. (A) Downregulation of *PLUTO* (HI-LNC71) and *PDX1* in islets from donors with T2D or IGT. Differential expression analysis was performed on control (n=50) versus T2D (n=10) or IGT (n=15) samples. Boxplots represent expression normalized to mean of control samples. Adjusted p-values are shown. (B) Schematic representation of the human *PDX1* locus and its associated enhancer cluster. 4C-seq analysis was designed to identify regions interacting with the *PDX1* promoter region in EndoC-βH1 cells. Red and orange vertical lines depict active and poised islet enhancers, respectively. F and R represent forward and reverse RNA-seq strands, scales represent RPM. *PLUTO (HI-LNC71)* was generated from a *de novo* assembly of islet RNA-seq, and differs from a transcript annotated in UCSC and RefSeq that originates from a *PDX1* intronic region. (C) Downregulation of *PLUTO* or *PDX1* using amiRNAs resulted in reduced *PDX1* mRNA and protein levels. EndoC-βH1 cells were transduced with control (black), *PLUTO* (white) or *PDX1* (turquoise) amiRNA vectors 80 hours prior to harvest. RNA levels were assessed by qPCR, normalized to *TBP* and expressed as fold over control amiRNA samples (n=4). For protein quantification, PDX1 levels were first normalized to the average of TBP and H3 levels and then compared to the control amiRNA sample. (D) Downregulation of *PLUTO* in human islet cells results in reduced *PDX1* mRNA levels. Islet cells were dispersed and transduced with amiRNA vectors (n=3) as in (B). (E) Downregulation of *PLUTO* in EndoCβH3 cells using CRISPRi also decreases *PDX1* mRNA. EndoC-βH3 cells were nucleofected with CRISPRi vectors 80 hours prior to harvest. RNA levels were assessed by qPCR, normalized to *TBP* and then to control CRISPRi sample (n=3). (F) *PDX1* and *PLUTO* RNA levels were highly correlated in 64 human islet samples. (G) Knockdown of *PDX1* and *PLUTO* resulted in differential expression of similar genes. Fold change value (Log2) of top 250 dysregulated genes following the *PDX1* knockdown was plotted against the same genes following the *PLUTO* knockdown. (H) Gene Set Enrichment analysis (GSEA) showed that genes that were downregulated upon knockdown of *PDX1* and *PLUTO* were enriched in genes whose enhancers were bound by PDX1 (red) in islets, but not in 10 control gene sets (black) that were expressed at similar levels as PDX1-bound genes. (I) Knockdown of *PDX1* and *PLUTO* resulted in differential expression of genes with similar biological process annotations. (J) Examples of known *PDX1* regulated genes that are also coregulated by *PLUTO* in parallel knockdown experiments. mRNA levels were assessed as in (B). Error bars denote SEM, *** p<10^−3^, **p<0.01, *p<0.05 (Student’s t test).

*PLUTO* is a multi-isoform transcript that contains five major exons that span nearly 100 kb, encompassing a cluster of enhancers that make three-dimensional (3D) contacts with the *PDX1* promoter in human islets and in EndoC-βH1 cells (**Figure 6B**, **S6A**). This observation suggested that *PLUTO* could affect cis-regulation of the *PDX1* gene.

To test whether *PLUTO* regulates *PDX1*, we first examined EndoC-βH1 cells after amiRNA-mediated knockdown of *PLUTO* RNA, and found reduced *PDX1* mRNA and protein levels (**Figure 6C**). Similarly, knockdown of *PLUTO* RNA in dispersed primary human islet cells caused decreased *PDX1* mRNA (**Figure 6D**). To validate these experiments through a complementary approach, we used CRISPR interference (CRISPRi), which involves targeting guide RNAs (gRNAs) downstream of a gene’s transcriptional initiation site to block its transcription. Two independent gRNAs that targeted a region downstream of the *PLUTO* initiation site efficiently reduced *PLUTO* RNA levels relative to non-targeting gRNAs, and in both cases this led to decreased *PDX1* mRNA expression (**Figure 6E**). Therefore, perturbing either *PLUTO* RNA levels or its transcription leads to the same inhibitory effect on *PDX1* mRNA.

The mouse *Pdx1* locus also has an islet lncRNA *(Pluto)* that shows only limited sequence homology with human *PLUTO. Pluto* is also transcribed from the opposite strand of *Pdx1*, but is initiated from a promoter within the first intron of *Pdx1*, and like *PLUTO*, spans a broad regulatory domain upstream of *Pdx1* (**Figure S6C**). Knockdown of *Pluto* RNA in the mouse β cell line MIN6 also led to decreased *Pdx1* mRNA levels (**Figure S6E**). These experiments therefore indicated that *PLUTO* regulates *PDX1* mRNA in human β cell lines and primary islet cells, and an analogous effect was observed for the mouse lncRNA ortholog.

Consistent with this regulatory relationship, *PLUTO* and *PDX1* RNA levels are highly correlated across islet samples (Pearson’s r=0.86, p=10^−15^, **Figure 6F**), and knockdown of *PDX1* and *PLUTO* in EndoC-βH1 cells resulted in the deregulation of a shared set of genes (**Figure 6G-J**). Furthermore, *Pluto* and *Pdx1* were found to be regulated with nearly identical dynamics in response to a shift in glucose concentration (4 to 11 mM) in mouse pancreatic islets (**Figure S6D**). *PLUTO* and *PDX1* therefore regulate a common program in pancreatic islets, and this is at least in part explained by the fact that *PLUTO* regulates *PDX1*.

### *PLUTO* regulates *PDX1* transcription and local 3D chromatin structure

To assess the mechanisms underlying the function of *PLUTO*, we first examined if *PLUTO* controls the stability or transcription of *PDX1.* Transcriptional inhibition experiments using Actinomycin D showed no significant differences in the stability of *PDX1* mRNA upon *PLUTO* knockdown (**Figure 7A**). By contrast, intronic *PDX1* RNA was reduced upon *PLUTO* knockdown, suggesting that *PLUTO* regulates *PDX1* transcription (**Figure 7B**).

**Figure 7.**
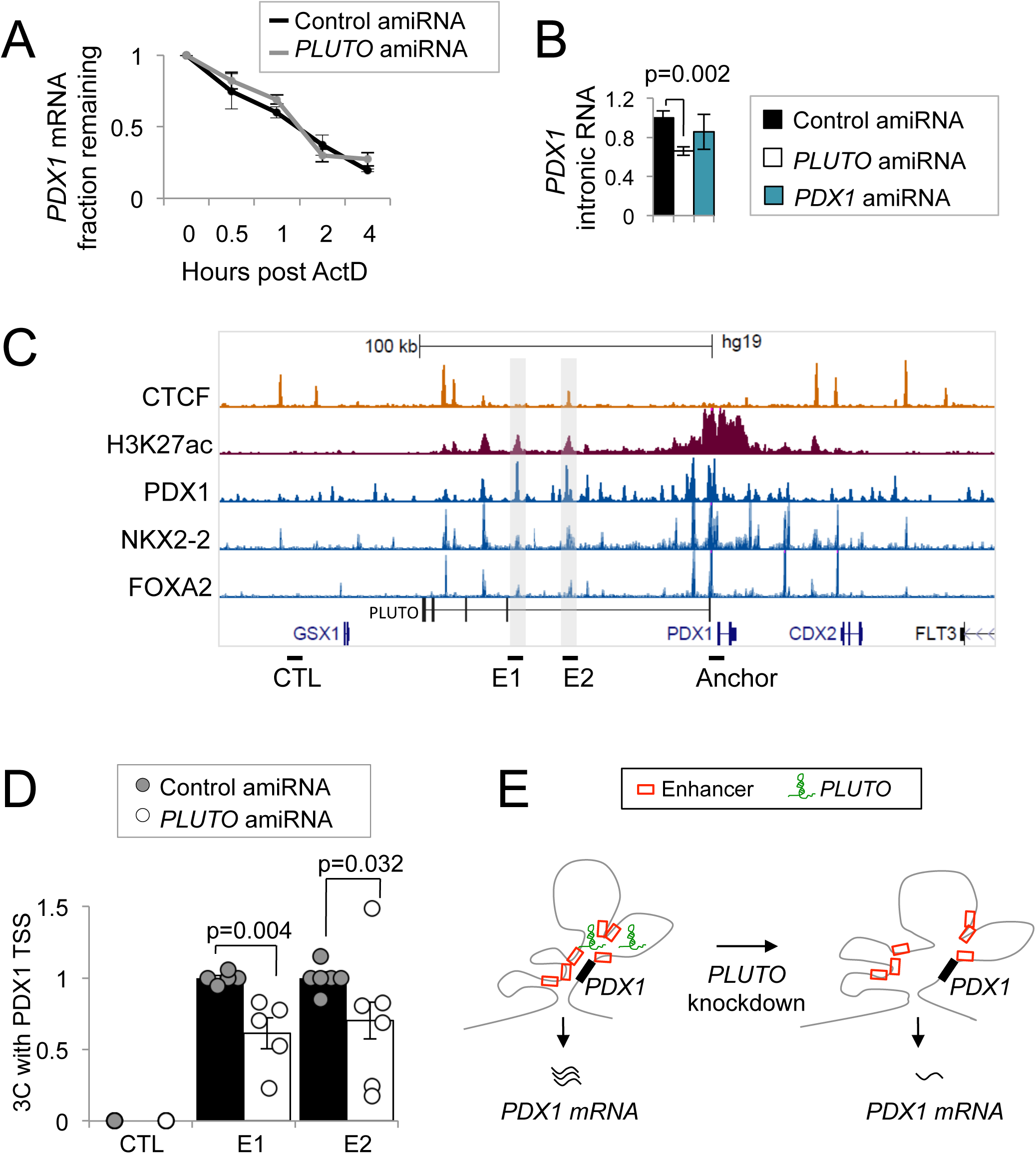
*PLUTO* regulates *PDX1* transcription and 3D chromatin structure. (A) The mRNA stability of *PDX1* was unaffected by *PLUTO* knockdown. *PDX1* mRNA was measured in control and *PLUTO* amiRNA knockdown in EndoC-βH1 cells after Actinomycin D (ActD) treatment (n=3). mRNA levels are presented as a percentage of levels observed at time=0. (B) Knockdown of *PLUTO* was carried out as in **Figure 6B**, and this led to reduced *PDX1* transcription as assessed by qPCR analysis of intronic *PDX1* RNA levels using hydrolysis probes. Values were normalized to *TBP* mRNA and expressed as fold over control amiRNA sample (n=4). (C) Schematic of selected epigenomic features of the *PDX1* locus, (D) *PLUTO* is required for 3D contacts between the *PDX1* promoter and distal enhancers. 3C analysis revealed that knockdown of *PLUTO* resulted in reduced contacts between *PDX1* promoter (anchor) and two enhancers (E1,E2). Interaction signals were normalized to a control region on *PDX1* intron. CTL represents a negative control region that does not harbor interactions with the *PDX1* promoter. Error bars denote ±SEM, p values are from a Student’s t test. (E) *PLUTO* knockdown resulted in impaired 3D contacts between the *PDX1* promoter and its adjacent enhancer cluster, causing reduced *PDX1* transcriptional activity.

Because *PLUTO* spans an enhancer cluster, we hypothesized that it could regulate the chromatin state of active enhancers. We thus knocked down *PLUTO* in β cells and measured H3K27 acetylation, as well as H3K4 mono and tri-methylation levels at several enhancers within the cluster. Our results indicate no significant changes in these characteristic active chromatin marks (**Figure S7**).

We next determined whether *PLUTO* affects the 3D contacts between the enhancer cluster and the *PDX1* promoter. Examination of the *PDX1* locus using quantitative chromatin conformation capture (3C) assays revealed that two far upstream enhancers (**Figure 7C**) showed reduced contacts with the *PDX1* promoter after *PLUTO* knockdown (**Figure 7D**). These findings therefore show that *PLUTO* regulates the transcription of *PDX1*, a key pancreatic β cell transcriptional regulator, and that this is associated with its ability to promote contacts between the *PDX1* promoter and its enhancer cluster (**Figure 7E**).

## Discussion

In the current study we have tested the hypothesis that lncRNAs play a role in cell-specific gene regulation in pancreatic β cells, a cell type that is central in the pathogenesis of human diabetes. We have thus carried out for the first time a systematic analysis of the function of a set of human β cell-specific lncRNAs. Our experiments revealed several examples of β cell lncRNAs in which sequence-specific perturbation causes transcriptional and functional phenotypes. We have further shown that β cell-specific lncRNAs and TFs regulate a common transcriptional network. Finally, we have demonstrated that β cell-specific lncRNAs directly or indirectly participate in the regulation of human enhancer clusters, which are the major functional targets of islet-specific transcription factors and key cis-regulatory determinants of islet cell transcriptional programs (Pasquali et al., 2014). Importantly, these conclusions are supported by concordant results from co-expression network analysis and loss of function experiments. These studies should be interpreted in light of previous evidence indicating that a significant fraction of lncRNAs show lineage-specific expression (Cabili et al., 2011; Derrien et al., 2012; Goff et al., 2015; Guttman et al., 2011; Iyer et al., 2015; Moran et al., 2012; Pauli et al., 2012). Our study extends previous findings by demonstrating a functional role of lncRNAs in lineage-specific TF networks.

Our findings invite the question of what molecular mechanisms underlie the regulatory effects of β cell lncRNAs. LncRNAs have been proposed to control gene expression through diverse molecular mechanisms, including the formation of protein-specific interactions and scaffolds, RNA-DNA or RNA-RNA hybrids, the titration of miRNAs, and the modulation of 3D chromosomal structures (Rinn and Chang, 2012; Wang and Chang, 2011), while some transcripts currently defined as lncRNAs can theoretically encode for atypical small peptide sequences (Andrews and Rothnagel, 2014). Our knockdown and co-expression analyses have identified a subset of functional lncRNAs that appear to regulate a nearby gene, suggesting a lncRNA-based cis-regulatory mechanism, while others are likely to exert transregulatory effects. We focused on one functional nuclear-enriched β cell lncRNA, *PLUTO*, and found that its function in β cell networks is at least in part due to its ability to elicit an effect on the transcription of its adjacent gene, *PDX1*, which encodes for a key β cell transcription factor. Importantly, this was observed for both the mouse and human orthologs, and similar effects were obtained through RNAi suppression or through CRISPR–induced transcriptional interference of *PLUTO.* Our studies further showed that *PLUTO* promotes 3D interactions between the *PDX1* promoter and its upstream enhancer cluster, which is contained within the body of the *PLUTO* gene. We thus propose that *PLUTO* regulates the 3D architecture of the enhancer cluster at the *PDX1* locus. This finding is reminiscent, yet distinct from earlier examples of non-coding RNA genes that modulate 3D chromosomal structure (Lai et al., 2013; Yao et al., 2010). Given that a significant number of lncRNAs are co-expressed with adjacent lineage-specific protein-coding genes, it is possible that the general regulatory paradigm described here is relevant to analogous lncRNA-protein coding gene pairs.

Taken together, our data implicate cell-specific lncRNAs in human β cell transcriptional programs. Given the importance of TFs in the pathophysiology of human diabetes and their role in β cell programming strategies, it now seems reasonable to explore whether β cell lncRNAs also play analogous roles (Bell and Polonsky, 2001; Flanagan et al., 2014; Zhou et al., 2008). The findings reported here therefore strengthen earlier suggestions that defects in β cell lncRNAs might contribute to the pathogenesis of human diabetes (Fadista et al., 2014; Moran et al., 2012), and warrant an assessment of whether they can be harnessed to promote β cell differentiation, function or cellular mass.

## Experimental procedures

### Pancreatic Islets

Human islets used for RNA-seq and ChIP-seq were cultured with CMRL 1066 medium containing 10% Fetal Calf Serum (FCS) before shipment, after which they were cultured for three days with RPMI 1640 medium containing 11 mM glucose, supplemented with 10% FCS.

### Glucose stimulated insulin release

Glucose stimulated insulin release was assayed in EndoC-βH1 or EndoC-βH3 cells as described (Benzara et al., 2015, Ravassard et al., 2011).

### RNA analysis

RNA was isolated with Tripure (Roche) and treated with DNase I (Sigma). qPCR was performed with SYBR green or Taqman probe detection (van Arensbergen et al., 2010). See **Table S4** for oligonucleotide and probe sequences.

### amiRNA and CRISPRi experiments

Lentiviral vectors carrying amiRNAs targeting TFs, lncRNAs and non-targeting control sequences were transduced into the EndoC-βH1 human β cell line as described (Castaing et al., 2005; Ravassard et al., 2011; Scharfmann et al., 2014). **Figure S1A** illustrates the vector design. Oligonucleotide sequences are shown in **Table S4**. Non-transduced cells were assayed in parallel. Cells were harvested at 80 hours post transduction for RNA extraction. For transduction of human islets, islets were first dispersed using trypsin-EDTA and gentle agitation. CRISPRi experiments were performed with two gRNAs designed to target *PLUTO* exon 1, or two unrelated intergenic control regions, and transfected in EndoC-βH3 cells (**Table S4**).

### Gene expression array analysis

RNA was hybridized onto HTA2.0 Affymetrix arrays. RMA normalization was carried out using Expression Console (Affymetrix). Gene based differential expression analysis was done using Transcriptome Analysis Console (TAC, Affymetrix). Enhancer cluster genes were defined by genes that were associated with clustered islet enhancers that show top 50 percentile binding by TFs (PDX1, FOXA2, NKX2-2, NKX6.1, MAFB), as defined previously (Pasquali et al., 2014). Pancreatic islet gene sets used for enrichment analysis are shown in **Table S5**. A list of islet-enriched genes was generated as those with more than two standard deviations higher expression in human islets than the average expression in 16 human tissues (**Table S5**). Data (cel and chp files) can be found at Gene Expression Omnibus (GEO, accession: GSE83619).

### Differential expression in IGT and T2D islets

RNA-seq data has been previously described (Fadista et al., 2014). The samples were aligned to the hg19 genome using STAR aligner version 2.3.0 as described in supplemental methods, quantification was carried out with HTseq-Count 0.6.1, and differential expression analysis of lncRNA genes was done using DEseq2 1.10 (**Table S3**), using an adjusted p-value threshold of 0.05.

### Chromatin conformation capture (3C)

3C and 4C-seq was carried out as previously described (Pasquali et al., 2014; Tena et al., 2011) For real-time PCR quantification, readings were normalized to a control region within the *PDX1* intron. Normalized values are expressed as a fraction of nontargeting amiRNA control sample. See **Table S4** for oligonucleotide sequences.

### Annotation of islet lncRNAs

LncRNAs were annotated through de novo assembly of ∼5 billion stranded paired-end RNA-seq reads from 41 human islet samples, filtered for expression in FACS-purified β cell cells, lack of enrichment in the pancreatic exocrine fraction to exclude acinar contaminants, and presence of presence of H3K4me3 enrichment in the vicinity of the 5’ end. A more detailed description of the annotation process is provided in supplemental methods. Annotations are available in **Table S3** and can be accessed on a UCSC genome browser (GRCh37/hg19) session by selecting “track hubs”, and selecting “Human Islet lncRNAs”. Alternatively the track hub can be directly visualized in the <u>UCSC Genome Browser</u>.

### Network analysis

WGCNA(v2) tool was used to build a co-transcriptional network based on mRNAs from 64 human islet RNA-seq samples.

## Author contributions

J.F. and I.A. conceived the idea, designed experiments and wrote the manuscript. J.F. supervised and I.A. coordinated the project. I.A., Z.T., H.W., J.Y., C.A., E.S., A.S., L. Pasquali and D.M.Y.R. contributed to data analysis. I.A., A.B., M.B., C.S.C., R.G.F., J.G.H. and N.C. performed experiments. D.M.Y.R, I.M. and N.N. annotated lncRNAs. L.Piemonti, T.B., C.B., J.K.C., F.P. provided samples. I.A., J.F., Z.T., A.B., D.M.Y.R, L.G., C.B., J.K.C., F.P., P.R., A.S., L.G., C.A., E.S. discussed results. All authors read and approved the manuscript. Z.T, A.B., D.M.Y.R, C.S-C. contributed equally.

## Acknowledgements

This research was supported by the National Institute for Health Research (NIHR) Imperial Biomedical Research Centre. Work was funded by grants from the Wellcome Trust (WT101033 to J.F.), NIH-BCBC (2U01 DK072473-06 to J.F., P.R) Ministerio de Economía y Competitividad (BFU2014-54284-R to J.F.) and Horizon 2020 (667191 to J.F.). Work in IDIBAPS was supported by the CERCA Programme, Generalitat de Catalunya. J.Y. was supported through Berg and Unity Biotechnology fellowship. The authors are grateful to Helena Raurell Vila for experimental help and Romain Derelle and Loris Mularoni for advice in bioinformatic analysis. P. R. is a shareholder and consultant for Endocells/Unicercell Biosolutions. Z.T. receives financial support from Berg Pharma and Unity Biotechnology as a consultant.

